# Treatment with a polyherbal extract improves fat metabolism, attenuates hepatic stellate cell activation and fibrogenesis

**DOI:** 10.1101/2021.08.06.455195

**Authors:** KB Pavan, Mirza Rizwan Baig, Mallappa Onkar Murthy, Mohammed Azeemuddin, V.R Hariprasad, Mohamed Rafiq, Raghavendra P Rao

## Abstract

Non-alcoholic steatohepatitis (NASH) involves dysregulations in *denovo* lipogenesis, fatty acid oxidation, and fibrogenesis. Targeting these pathways holds promise for the treatment of liver disorders. Here we test the extract of a polyherbal formulation (namely Liv.52), which is approved by the Government of India’s Drug Regulatory Authority - AYUSH. The current study evaluates the effect of Liv.52 on *denovo* lipogenesis, fatty acid oxidation, and fibrogenesis. Both *in vivo* and *in vitro* model systems were employed to evaluate the efficacy of this polyherbal formulation. Male Wistar rats were dosed with Liv.52 for 2 weeks (250mg/k.g) and expression levels of the genes involved in *de novo* lipogenesis and fatty acid oxidation pathways were analysed by quantitative real time PCR. Liv.52 treatment resulted in increased hepatic fatty acid oxidation and decreased *de novo* lipogenesis in these rats. It also reduced hepatic stellate cell activation in CCL_4_ treated Wistar rats as evidenced by histological evaluation. For *in vitro* experiments, HepG2 cells were cultured under lipotoxic conditions (using 200μM palmitic acid) and the conditioned media from these cells were used for inducing activation and fibrogenesis in human hepatic stellate cells (HHSteC). Treatment with lipotoxic conditioned media resulted in activation of hepatic stellate cells and fibrogenesis, as evidenced by increased expression of α-smooth muscle actin (α-SMA), and desmin (markers of stellate cell activation) and increased levels of collagen and lumican (markers of fibrogenesis). Treatment with Liv.52 reversed the up-regulation of α-SMA, collagen and lumican levels in HHSteC cells. These results indicate that Liv.52 exerts its hepatoprotective effect by improving fatty acid metabolism and fibrogenesis.

## INTRODUCTION

Non-alcoholic fatty liver disease (NAFLD) has emerged as a leading chronic liver disorder in the developed countries and its prevalence is growing even in the developing countries[1]. Most of the patients with NAFLD exhibit simple fatty liver while 5 to 20 percent of patients with fatty liver progress to non-alcoholic steatohepatitis (NASH), of which 10–20% develop higher-grade fibrosis and <5% progress to full-blown cirrhosis[2]. The progression of NAFLD into NASH and subsequent fibrosis is a very complex process involving dysregulation of several cellular pathways such as *de novo* lipogenesis, fatty acid oxidation and stellate cell activation (discussed in the results and discussion section) and it is also the result of functional dysregulation of several cell types – hepatocytes, immune cells, fibroblasts and hepatic stellate cells[3]. Hence, an effective therapeutic strategy for treating the NASH and subsequent fibrosis should be aimed at correcting all these dysregulations. Current therapeutic strategies for NASH include insulin sensitizers, lipid-lowering drugs, anti-inflammatory drugs and anti-fibrotic agents[4]. While these drugs are effective in their respective actions, they may not address all features of NAFLD -from NASH to fibrosis. Recent advances in our understanding of systems biology suggest that polypharmacology approaches might be more effective for treating complex diseases rather than targeting only specific pathways and drug targets[5]. Therapeutic systems involving herbal formulations greatly rely on polypharmacology wherein the phytoactives target several cellular processes key to the disease progression. Several plant based natural products have been shown to be effective as hepatoprotectants and in the absence of an established treatment regime for NASH and liver fibrosis, much attention has been focussed on developing the natural product based formulations. Liv.52 is a polyherbal formulation approved by the Government of India’s Drug Regulatory Authority, the department of Ayurveda, Yoga, Naturopathy, Unani, Siddha and Homoeopathy (AYUSH) of the Ministry of Health and Family Welfare (New Delhi, India). Clinical and preclinical studies indicate its efficacy as a hepatoprotective agent in NASH and cirrhosis[6,7].

In the current study, we have evaluated the effects of Liv.52 for its effect on fat metabolism, hepatic stellate cell activation and fibrosis. We provide the evidence that Liv.52 reduces *de novo* lipid synthesis, improves fatty acid oxidation, increases mitochondrial function and inhibits stellate cell activation and extracellular matrix (ECM) accumulation, all of which are key events during the process of progression of NAFLD into NASH and subsequent fibrosis.

## RESULTS DISCUSSION AND CONCLUSION

In the current study we studied the effect of Liv.52 on

1. *de novo* lipogenesis –(using rat model)
2. Fatty acid oxidation – (using rat model)
3. Hepatic stellate cell activation and fibrogenesis – (using human hepatic stellate cells HHSteC) *De novo* lipogenesis and fatty acid oxidation are important determinants of steatosis[11–14], while stellate cell activation and fibrogenesis and subsequent processes which lead to fibrosis. In the following sections these results are describe in detail.

### I. Effect of Liv.52 on fat metabolism- *in vivo* studies

Two of the important factors that impact the lipid accumulation in the hepatocytes (steatosis) are

1. Increased denovo lipogenesis – Metabolic overload conditions such as increased carbohydrate intake, leads to conversion of excess glucose into triglycerides through the action of key enzymes involved in lipogenic and glycolytic pathway[11,12].
2. Impaired fatty acid oxidation - Apart from denovo lipogenesis, the other arm of metabolic pathway which contributes to steatosis, is impaired hepatic fatty acid oxidation[13]. Impaired fatty acid oxidation has been shown to result in steatosis[14] and enhancing the fatty acid oxidation is shown to decrease steatosis[15]. Besides, free fatty acids in the hepatocytes can cause lipotoxicity[16]. Hence handling these free fatty acids becomes very important in order to deal with the toxicity and this can be achieved by sequestering these fatty acids to beta oxidation pathway

With this background, we evaluated the effect of Liv.52 treatment on *de novo* lipogenesis and fatty acid oxidation.

#### Effect of Liv.52 on de novo lipogenesis

Wistar rats were dosed with Liv.52 (250mg/kg p.o) for 2 weeks and the liver samples were analysed for expression levels of key genes involved in *denovo* lipogenesis. As indicated in **Fig.1A** treatment with Liv.52 reduced the expression of ChREBP (Carbohydrate-responsive element-binding protein) to 60% of control. ChREBP is a glucose-dependent transcription factor and is a key regulator of hepatic lipogenesis, which can induce lipogenesis independent of insulin[17]. It upregulates hepatic lipogenesis by transcriptionally controlling key enzymes involved in glycolysis and fatty acid synthesis[18,19]. To further confirm this, we tested the expression levels of the downstream targets of ChREBP viz. glucokinase (a glycolytic enzyme) and ACC1 (Acetyl coA carboxylase-1). Glucokinase activity increases the generation of glycerol backbone necessary for the synthesis of triglyceride and the enzyme ACC1 catalyses the first step in the *de novo* biosynthesis of fatty acids. A coupled activity of these two enzymes can result in increased production of liver triglycerides[20]. Indeed, it was observed that Liv.52 treatment reduced the levels of these enzymes (**Fig.1A**) to 57% and 41% respectively. These studies indicate that Liv.52 reduces the *de novo* lipogenesis in liver by acting at the level of ChREBP. In the light of these observations, some the protective effects of Liv.52 on ethanol induced liver toxicity[21] could also be due to its ability to quench *de novo* lipogeneis. *De novo* lipogenesis is in fact one of the key mechanisms through which ethanol induces steatosis. Ethanol consumption leads to generation of cytosolic acetyl coenzyme A (acetyl-CoA) which serves as a substrate for fatty acid synthesis.

**Figure-1.**
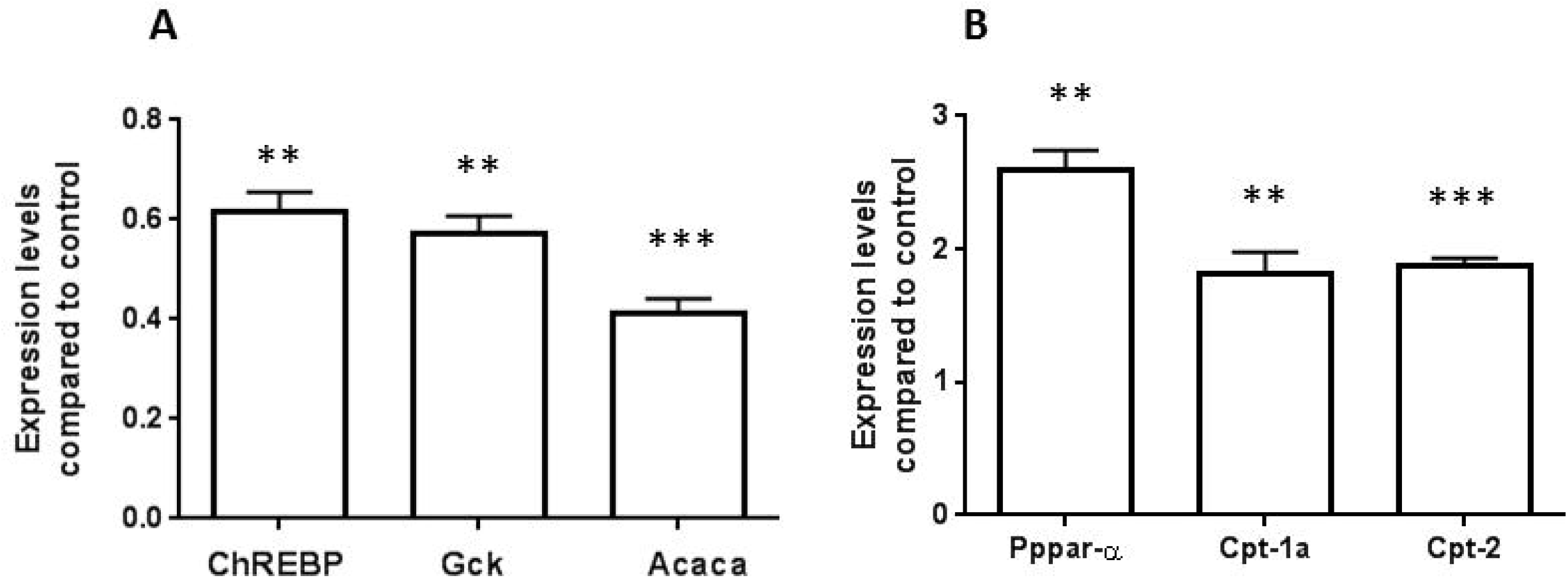
Effect of Liv.52 on denovolipogenesis and fatty acid oxidation : Total RNA was isolated from rat livers (n=6) and was converted into c-DNA (A & B). The expression level of indicated genes (both A & B) was estimated by using real time quantitative PCR. Expression levels of all the genes in control rats were normalized as 1 (both A & B). Two tailed unpaired t-test was used for statistical analysis. ** p<0.01, ***p<0.001 compared to respective controls

#### Effect of Liv.52 on fatty acid oxidation

The liver samples from the Liv.52 treated rats were analysed for levels of key genes involved in fatty acid oxidation. As indicated in **Fig.1B**, in our experiments we observed an increase in the expression levels of PPAR-α (2.6 fold), CPT-1a (1.8 fold), and CPT-2 (1.9 fold) in rats treated with Liv.52. The transcription factor PPAR-α controls the expression of several enzymes involved in beta oxidation of fatty acids in the liver[15]. CPT-1a and CPT-2 are involved in transfer of long chain fatty acyl CoA from cytosol to mitochondrial matrix. Earlier studies have indicated that a moderate increase in the CPT-1a activity is sufficient to reduce hepatic triglyceride accumulation[22] and interventions that increase the CPT-1 have been suggested as a means for treatment of NAFLD. Supporting the role of Liv.52 further in fatty acid oxidation, it was observed that Liv.52 treatment enhanced the activity of 3-hydroxy acyl coA dehydrogenase activity in HepG2 cells (**Fig.4**). 3-hydroxy acyl coA dehydrogenase is an enzyme involved in mitochondrial beta oxidation of fatty acids.

The above data taken together suggest that Liv.52 enhances fat oxidation in the liver while reducing the *de novo* lipogenesis.

### II. Effect of Liv.52 on fibrogenesis – in vitro studies

In a physiological set up, chronic metabolic overloading leads lipid accumulation in hepatocytes and subsequent injury[3]. This initial injury to the hepatocytes acts a trigger for several of cellular events in the liver which finally results in activation of quiescent hepatic stellate cells into myofibroblasts, which leads to excess production of ECM[23]. Earlier studies have indicated that treatment of hepatocytes with saturated fatty acids like palmitic acid can lead to accumulation of triglyceride, trigger inflammation and exhibit hallmarks of steatosis[3,9,24,25]. Hence this treatment is proposed as an *in vitro* model for steatosis. Studies also indicate that conditioned media obtained from hepatocytes cultured under high metabolic overload, can trigger hepatic stellate cell activation [26]. In our experiments we employed this as a model of steatosis *in vitro*. HepG2 cells when cultured in presence of palmitic acid for 24 hours, showed increased triglyceride levels as measured by quantification of oil-o-red staining (**Fig.2A**). During this period no appreciable cytotoxicity was seen in the cells (data not shown). The conditioned media was collected from these cells and processed as explained materials and methods. This conditioned media was referred to as lipotoxic conditioned media (LTCM). The conditioned media obtained from the cells grown without palmitate but only with BSA (which was used for conjugation of palmiate) was used as control conditioned media (CCM). The human hepatic stellate cells HHSteC were cultured with this conditioned media for 24 hours. Following this, the stellate cell activation and fibrogenesis were studied.

**Figure-2.**
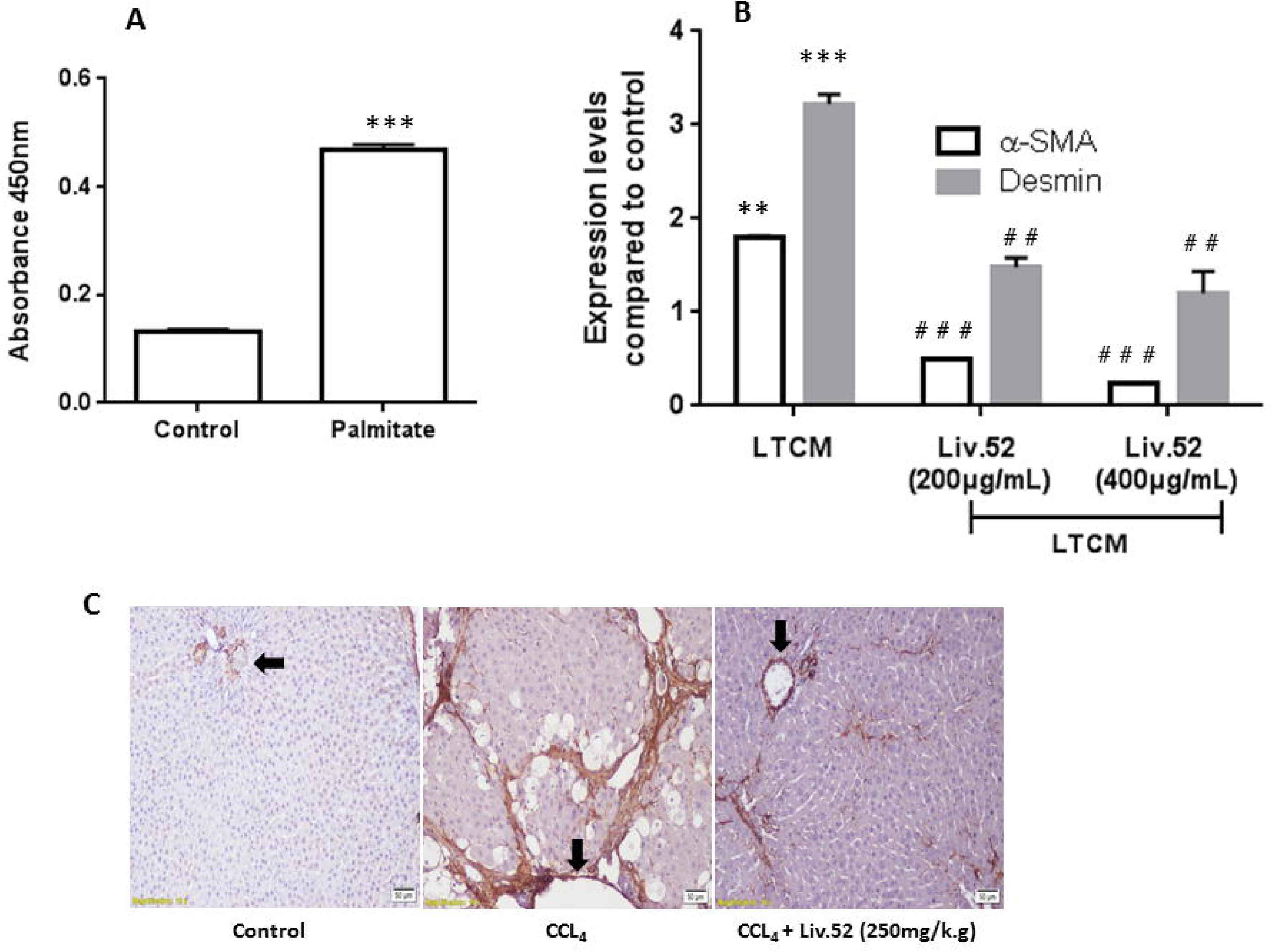
Effect of Liv.52 on hepatic stellate cell activation: **(A)** -HepG2 cells were cultured either in the absence (control) or presence of 200μM palmitate as indicated in materials and methods. The cells were stained with oil O-red and the dye was finally solubilized in 10%DMSO and quantified spectrophotometrically at 450nm. Two tailed unpaired t-test was used for statistical analysis. ** p<0.01, ***p<0.001 compared to respective controls. **(B)**-HHSteC cells were incubated with lipotoxic conditioned media (LTCM) obtained from HepG2 cells as explained in materials and methods and the gene expression levels were quantified by real time quantitative PCR. Expression levels of all the genes in control cells (cells grown under non lipotoxic conditions as explained in materials and methods) were normalized as 1. Two tailed unpaired t-test was used for statistical analysis. ** p<0.01, ***p<0.001 compared to respective controls, ##p<0.01, ###p<0.001 when compared to LTCM treatment **(C)**- Immunohistochemistry of liver sections indicating staining for α-SMA protein in control rats, rats treated with CCL4 and rats treated with CCL_4_+Liv.52 as indicated in materials and methods. Arrows mark the portal vein.

#### Effect of Liv.52 on hepatic stellate cell activation

Our results show that lipotoxic conditioned media treatment resulted in activation of stellate cells as measured by expression levels of and Desmin. α-SMA is an established marker for hepatic stellate cell activation and as indicated in **Fig.2B**, treatment with LTCM resulted in about 1.8 fold increase in the α-SMA levels while the treatment with Liv.52 very significantly reduced the levels (72% reduction with 200μg/mL and 87% reduction with 400μg/mL) well below even the control levels. Similar to its effect on α-SMA expression, Liv.52 partially restored the level of expression of desmin (**Fig.2B**) which was upregulated by 3.2 fold following treatment with LTCM. Desmin is an intermediate filament and is reported in several studies as a gold standard for stellate cell activation[3,27]. The inhibitory effect of Liv.52 on stellate cell activation was also confirmed even in a fibrotic rat model. When dosed with CCL_4_ (at the dose of 0.5ml/kg body weight, twice weekly for 11 weeks), rats exhibited increased serum SGOT and SGPT and histological hallmarks of fibrosis (data not shown). As indicated in **Fig.2C** Immunohistochemistry clearly showed an increased expression of α-SMA in CCL4 treated rats and there was a marked decrease in expression of this marker in the rats treated with Liv.52. These data taken together indicate that Liv.52 inhibits hepatic stellate cell activation.

#### Effect of Liv.52 on fibrogenesis

As indicated in **Fig.3A** and **3B** when treated with LTCM, the levels of collagen and lumican significantly increased in the human hepatic stellate cells. The collagen levels increased by about 38% when treated with LTCM. When Liv.52 was included during the course of treatment, the collagen levels reduced to the levels comparable to control levels (110% with 200μg/mL and 88% with 400μg/mL of Liv.52) indicating almost a complete reversal of the effect of LTCM. Lumican is an important ECM protein whose exaggerated secretion causes fibrosis. Following treatment with LTCM there was about 1.7 fold increase in the lumican levels in the hepatocytes and treatment with Liv.52 significantly reduced the lumican expression levels (**Fig.3B**).

**Figure-3.**
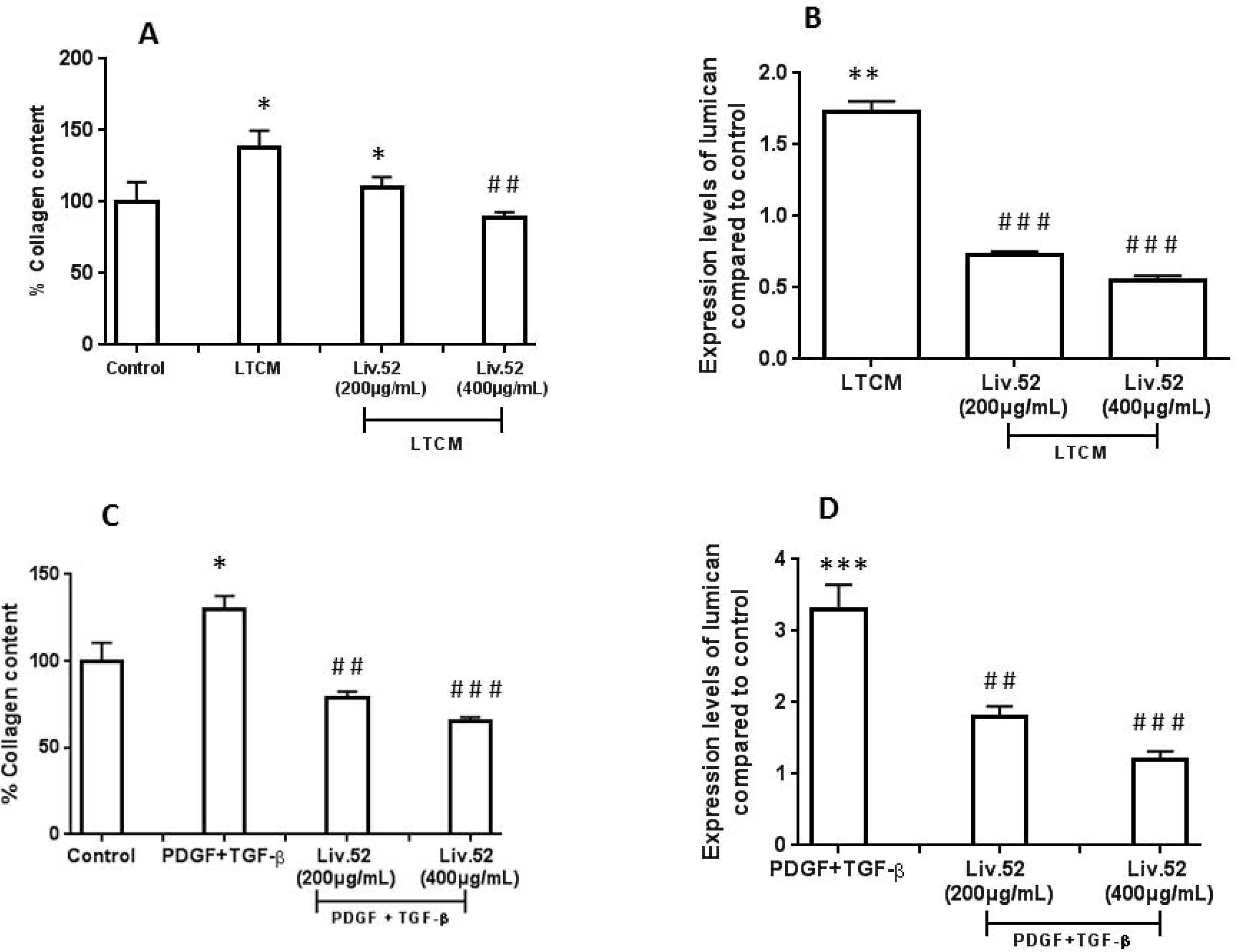
Effect of Liv.52 on fibrogenesis: Human hepatic stellate cells HHSteC were cultured either with the lipotoxic conditioned media (LTCM) obtained from HepG2 cells (A & B) or with 10ng/mL each of PDGF and TGF-β (C & D) as described in materials and methods. Collagen levels were estimated by sirus red staining (A & C) and lumican expression levels were estimated by quantitative real time PCR (B & D). For (B) & (D) the expression levels of lumican in the respective controls was normalized as 1. Two tailed unpaired t-test was used for statistical analysis. * p<0.05, ** p<0.01, p<0.001 compared to respective controls, ##p<0.01, ###p<0.001 when compared to LTCM or PDGF + TGF-β treatment

While hepatocyte injury and steatosis are the initial triggers for stellate cell activation, several other cell types and their cytokines also contribute to the process of stellate cell activation and fibrogenesis. Of these, TGF-β and PDGF are important cytokines that contribute at different stages of liver fibrosis such as stellate cell activation and fibrogenesis[28,29]. These cytokines are considered to be important for progression of fibrosis. TGF-β plays a major role in the transformation of hepatic stellate cells into myofibroblasts and stimulates the synthesis of extracellular matrix proteins while inhibiting their degradation. PDGF, on the other hand is a potent proliferative factor for hepatic stellate cells and myofibroblasts during liver fibrogenesis. In our experiments, we found that when incubated with 10ng/mL each of TGF-β β and PDGF cockatail, there was about 30% increase in the levels of cellular collagen and (**Fig.3C**) in the hepatic stellate cells. Collagen levels got reduced to 79% and 65% with 200μg/mL and 400μg/mL of Liv.52 respectively. (**Fig.3C**). Similarly, upon treatment with TGF-β β and PDGF cocktail, lumican levels increased to 3.3 fold further confirming the effect of the cocktail on fibrogenesis. Liv.52 effectively mitigated these effects and as indicated in **Fig.3D**, the lumican levels were reduced to 1.8 fold and 1.2 fold with 200μg/mL and 400μg/mL Liv.52 treatment respectively (**Fig.3D**).

#### Effect of Liv.52 on mitochondrial function

Mitochondrial dysfunction is proposed as a cause, effect or a concomitant feature in the development of NAFLD[13]. The mitochondrial function can be measured by activity of the enzyme 3-hydroxy acyl coA dehydrogenase. As indicated in **Fig.4** when HepG2 cells were treated with palmitate the activity of the mitochondrial enzyme was reduced by 30%. Upon treatment with, Liv.52 there was 20% (at 200μg/mL) and 50% increase (at 400μg/mL), in the enzyme activity. 3-hydroxyacyl coA dehydrogenase is an enzyme involved in mitochondrial fatty acid oxidation and these results further support our observations in the rat liver, in which there was an increase in the fatty acid oxidation pathway genes following treatment with Liv.52 (**Fig.1B**)

**Figure-4.**
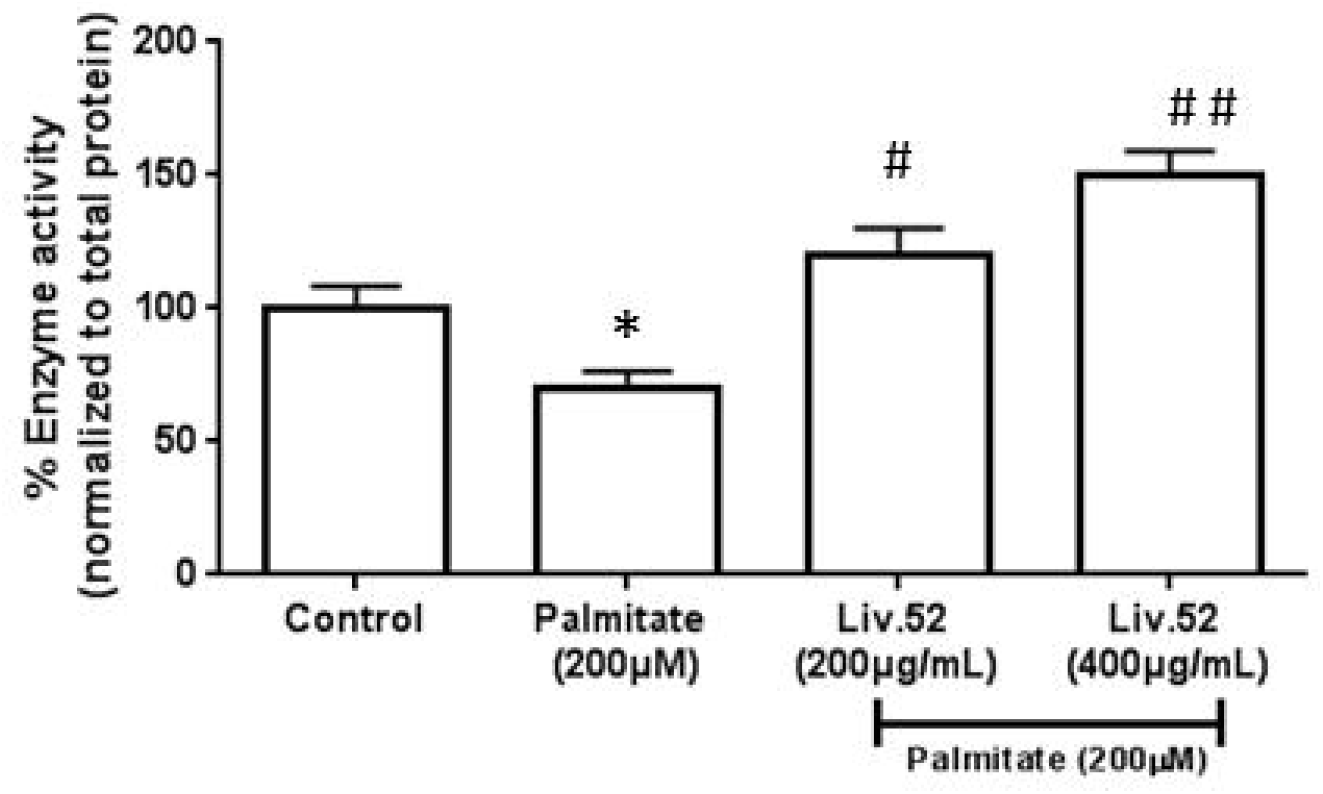
Effect of Liv.52 on mitochondrial function: HepG2 cells were cultured either in the absence (control) or presence of 200μM palmitate with or without Liv.52. The cells were lysed and the 3-hydroxyacyl CoA dehydrogenase activity was measured in these cells as described in materials and methods. The enzyme activity was normalized to total protein content in the cell lysate.

## Conclusions

Central to the process of development of progression of NAFLD into NASH is metabolic dysregulations which affect the balance between the ability of the liver to handle metabolites and the metabolite flux in the body. Progression of NASH into fibrosis, on the other hand involves several cell types such as hepatocytes, hepatic stellate cells and immune cells. Hepatic stellate cell activation and increased ECM production are key events during the progression of NASH into full-blown fibrosis. The pathology of NAFLD and its subsequent progression being a complex scenario, the treatment regime should involve therapeutic corrections at different levels. Our studies indicate that Liv.52, a polyherbal formulation-

- Reduces *de novo* lipogenesis
- Enhances mitochondrial fatty acid oxidation
- Inhibits stellate cell activation
- Inhibits accumulation of ECM proteins such as collagen and lumican.

Earlier studies have established the effect of Liv.52 in NASH and cirrhosis[6,7] and our studies demonstrate that Liv.52 exerts its effect against NASH by influencing aforementioned pathways. Taken together our study establishes the mechanism behind anti-NASH and anti-fibrotic effects of Liv.52.

## Materials and methods

### Liv.52 extract

The Liv.52 extract constitutes a mixture of *Capparis spinosa* (roots), *Cichorium intybus* (seeds), *Solanum nigrum* (whole plant), *Terminalia arjuna* (bark), *Cassia occidentalis* (seeds), *Achillea millefolium* (aerial part), and *Tamarix gallica* (whole plant). Preparation of Liv.52 extract and its detailed LC-MS/MS analysis are described earlier[6,8]. Briefly, the crude herbal materials were pulverized and the coarse powders obtained were blended. They were extracted with water and filtered through muslin cloth and concentrated in the reactors to attain 30% total solids. The soft extract was then subjected to spray drying to obtain the dry extract powder. At all stages, GMP (good manufacturing practice) and GLP (good laboratory practice) standards were maintained.

### Animal handling

Male Wistar rats were used for the experiments and they were housed in standard conditions of temperature (22 ± 3°C), relative humidity (55 ± 5%) and light (12 h light/dark cycle). Animals were fed with standard pellet diet (Provimi Animal Nutrition India Pvt. Ltd), and water ad libitum. The experimental protocols were approved by the Institutional Animal Ethics Committee (IAEC) of The Himalaya Drug Company, Bangalore, and the animals received humane care as per the guidelines prescribed by the Committee for the Purpose of Control and Supervision on Experiments on Animals (CPCSEA), The Ministry of Environment & Forests, Government of India.

For the experiments, male rats (8 week old) were divided into 2 groups of 6 each. Group 1 served as normal control and was administered with 10 ml/kg of water (p.o). Group 2 was treated with Liv.52 at a dose of 250 mg/kg (p.o). Dose of Liv.52 was selected based on the earlier studies and published literature [6]. All animals received the respective treatment for 2 weeks. 24 hours after the last dose of treatments, animals were euthanized and livers were excised and stored in Trizol.

For experiments with fibrotic model, twenty four male Wistar rats (12 weeks) were divided into 3 groups of 8 each based on the body weight. Group I rats were served as normal control, group II and III received carbon tetrachloride [0.5ml/kg i.p (intaperitoneal)] (which was suspended in olive oil 1:9 v/v) twice weekly for 11 weeks. The rats from groups I and II received water once daily at a dose of 10 ml/kg (p.o, per oral.) and served as normal and pathological controls respectively. Rats of group III received Liv.52 at a dose of 250 mg/kg body weight. All animals received the respective treatments for 11 weeks. At the end of 11 weeks serum levels of SGOT and SGPT were estimated by autoanalyser. Twenty four hour after the last dose treatment, the animals were euthanized. Livers were excised and fixed in 10% neutral buffered formalin. Immunohistochemistry was performed on 4-5μ thick paraffin embedded liver tissue sections mounted on Super frost positively charged microscopic slides (Fisher Scientific, USA) using the ready to use mouse monoclonal antibody (clone 1A4; Biogenex, Fremont, CA, USA) with advanced Super Sensitive Polymer-HRP IHC Detection System/DAB kit according to the manufacturer’s instructions (Biogenex, Fremont, CA, USA).

### Cell culture

HepG2 cells were routinely cultured in DMEM (Dulbecco’s Modified Eagle’s Medium) HG (high glucose) supplemented with10% FBS (fetal bovine serum) unless otherwise mentioned. Human hepatic stellate cells, HHSteC were routinely cultured in Stellate cell medium supplemented with 10% FBS.

### Preparation of palmitate

BSA (Bovine serum albumin) conjugated palmitate solution was prepared as described earlier [9]. Briefly, sodium palmitate (Sigma Aldrich) was dissolved at 500 mM concentration in 50% ethanol solution. This was further conjugated to BSA and 5 mM palmitate stock solution was obtained. This solution was filter sterilized and was used as stock solution.

### HepG2 Conditioned media preparation

HepG2 cells were grown to 80% confluence in DMEM HG supplemented with10% FBS in a humidified CO_2_ incubator at 37°C, after which the cells were serum starved for 8 hours. Following this, the cells were treated with lipotoxic media (DMEM containing 2% FBS and 200 μM BSA conjugated palmitate) for 24 hours and the media was collected and filtered through 0.2μ filter. This was referred to as lipotoxic conditioned media (LTCM) and was stored at −20°C until further use.

### Treatment of Stellate cells

Human primary heatpic stellate cells- HHSteC were seeded at a density of 10000 cells/ well in 96 well culture plate and allowed to reach 80 % confluence in HHSteC media. Cells were further washed thoroughly with cold PBS (phosphate buffered saline, pH 7.4), and the conditioned media obtained from HepG2 was added to the cells and the incubated for 24 hours with or without Liv.52. These cells were incubated for 24 hours. Appropriate vehicle controls were included. Similarly for experiments involving TGF-β and PDGF, the HHSteC cells were incubated with 10ng/ml each of these cytokines for 24 hours with or without Liv.52.

### Estimation of collagen

HHSteC cells were washed with PBS and cells were fixed in 10% formaldehyde for 30 minutes followed by PBS wash. 75μl of 1.2% sirus red reagent was added to each well and the plates were incubated at room temperature for 2 hours. The cells were later washed three times with 0.1N hydrochloric acid and the dye was finally extracted from each well into 75μl of 0.2N sodium hydroxide. The absorbance at 540nm was recorded for quantification. The collagen levels were calculated using the standard curve where pure collagen was used as standard.

### Quantitative real time PCR

The total RNA was isolated from the cells or rat liver using RNA isolation kit (Krishgen Biosystems, India). The quality of the RNA was confirmed on formaldehyde agarose gel with 1% strength by visualizing 28s and 18s bands. RNA was converted into cDNA using random hexamers as primers provided with the cDNA synthesis kit (Thermo Scientific, USA). cDNA was further subjected to quantitative real time PCR in CFX96 real-time system (Bio Rad, USA). The qPCR reaction contained cDNA, gene specific forward and reverse primers, 1X Syber green master mix (Bio Rad, USA) in a total reaction volume of 20 μl. The qPCR cycle comprised of 95 °C for 5 min, and 35 cycles of 95 °C for 30 s, annealing for 30 s, and 72 °C for 20 s. Melting curve profiles were analysed to confirm the amplification of the gene products. The expression levels of each gene were calculated by ddCt method using GAPDH as the house keeping gene.

#### β- hydroxyl acyl CoA dehydrogenase activity

HepG2 cells were seeded in 6 well plate at a density of 0.2million cells/well. Cells were grown until they reached 80% confluence in the presence or absence of Liv.52 for 24 hour incubation at 37°C. After 24 h incubation, supernatants were discarded and cells were lysed with lysis buffer containing protease inhibitor cocktail. Cell lysates were centrifuged at 1000xg for 10 min; the clear supernatant was used for estimating the enzyme activity [10]. The enzyme activity was normalized to protein levels.

## ACKNOWLEDGMENTS

Authors sincerely acknowledge Research and Development department of The Himalya Drug Company for supporting this study. We sincerely thank Dr.Gopumadhavan for his critical comments and suggestions which helped in improving the quality of the manuscript.

## COMPLIANCE WITH ETHICAL STANDARDS

This study does not involve human subjects. All applicable international, national, and/or institutional guidelines for the care and use of animals were followed. All procedures performed in studies involving animals were in accordance with the ethical standards of the institution or practice at which the studies were conducted.

